# Large and persistent social risks to forest carbon projects

**DOI:** 10.64898/2026.10.05.756721

**Authors:** Jacob I. Levine, Libby Blanchard, Danny Cullenward, William R.L. Anderegg

## Abstract

Despite the popularity of carbon credits as a tool to offset greenhouse gas emissions, there are growing concerns over whether their climate mitigation claims are robust. Most attention has focused on natural risks to forest carbon projects, such as wildfire, while social risks (e.g., those driven by fraud, mismanagement, and land conflicts) remain largely unquantified. Here we analyze 244 forest-based carbon projects to assess the vulnerability of the voluntary carbon market to social risks and find that projects systematically underestimate their risk of failure. Even under optimistic assumptions, projects are roughly twice as likely to fail within 25 years as their self-assessed risk implies over a 100-year horizon. Projecting these trends forward, we show that credits issued to failed projects could exceed the reserves registries hold to mitigate these risks within 1-2 decades. Overall, our results suggest stronger mechanisms and reserves are needed to ensure carbon credits deliver the promised climate benefits.

## Introduction

Over the last three decades, carbon credits have emerged as an increasingly popular tool to facilitate climate change mitigation (Kreibich, 2024; Ruiting et al., 2025; Trouwloon et al., 2023). In particular, the voluntary carbon market (VCM) has grown rapidly (Ruiting et al., 2025). Unlike compliance carbon markets, in which regulated entities surrender allowances or eligible carbon credits to meet legal obligations, the VCM involves corporations and individuals voluntarily purchasing carbon credits to meet self-determined climate commitments (Trouwloon et al., 2023). However, significant concerns have been raised regarding whether VCM credits represent genuine and durable emissions reductions or removals, calling into question voluntary carbon credits’ effectiveness as a climate mitigation tool (Anderegg et al., 2025; Haya et al., 2023; Probst et al., 2024; Romm et al., 2025; Tang et al., 2025; West et al., 2023).

A primary concern is the durability of carbon credits’ climate mitigation outcomes, particularly those that originate from nature-based carbon projects (Anderegg et al., 2025; Badgley et al., 2022; Haya et al., 2023; Marland et al., 2001; Wu et al., 2026). To fully offset the impacts of fossil CO_2_ emissions, a substantial fraction of which persist in the atmosphere for tens to hundreds of thousands of years, credited carbon must be sequestered for similar periods (Allen et al., 2022, 2025; Marland et al., 2001). Some registries have adopted a 100-year accounting standard for assessing durability risks (Anderegg et al 2025), but a period of at least 1000 years is needed to counteract the physical warming effects of CO_2_ emissions (Brunner et al., 2024). Although this discrepancy suggests VCM forest credits lack a one-to-one physical correspondence with fossil CO_2_ emissions, even 100-year durability guarantees are uncertain (Anderegg et al., 2025; Badgley et al., 2022; Cullenward et al., 2023; Haya et al., 2023; Wu et al., 2026).

The predominant threats to the durability of forest carbon storage include natural disturbances as well as sociopolitical, financial, and management risks (hereafter ‘social risks’). While a recent literature assesses natural risks (Anderegg et al., 2025; Badgley et al., 2022; Haya et al., 2023; Wu et al., 2026), very little is known about the adequacy of social risk estimates used in the VCM (Michaelowa et al., 2025).

Here we explore the vulnerability of forest-based VCM projects to social risks, using Verra—the largest voluntary registry, which issues two out of every three VCM carbon credits—as a case study. First, we conduct a systematic review of forest-based projects registered under Verra’s Verified Carbon Standard (VCS) to identify common risks and characterize the Verified Carbon Units (VCUs) it issues. We then forecast the potential impacts of social risks on VCU durability over the next several decades based on existing projects’ performance over 2000-2026.

### The buffer pool

Most carbon crediting mechanisms, including VCS, use a buffer pool to back nature-based carbon credits. Under this insurance-like system, projects contribute a portion of their credit issuance to a shared account to mitigate the loss of credited carbon storage, known as a reversal. When a project experiences a reversal due to factors like wildfire or illegal logging, it reports verified calculations of its losses to the registry, which cancels an equivalent number of credits in its buffer pool. This ensures that credits previously issued to that project still correspond to sequestered carbon and demonstrates that the underlying climate mitigation claim is robust.

Buffer pool operation depends on several related concepts (summarized in Table 1), including how long projects must be monitored to detect reversals and which reversal events the buffer pool covers. To determine each project’s required contribution to the buffer pool, carbon crediting mechanisms typically require a reversal risk assessment, which quantifies risks over a given time period (which we call the reversal risk assessment horizon, 100 years in VCS).

**Table 1:**
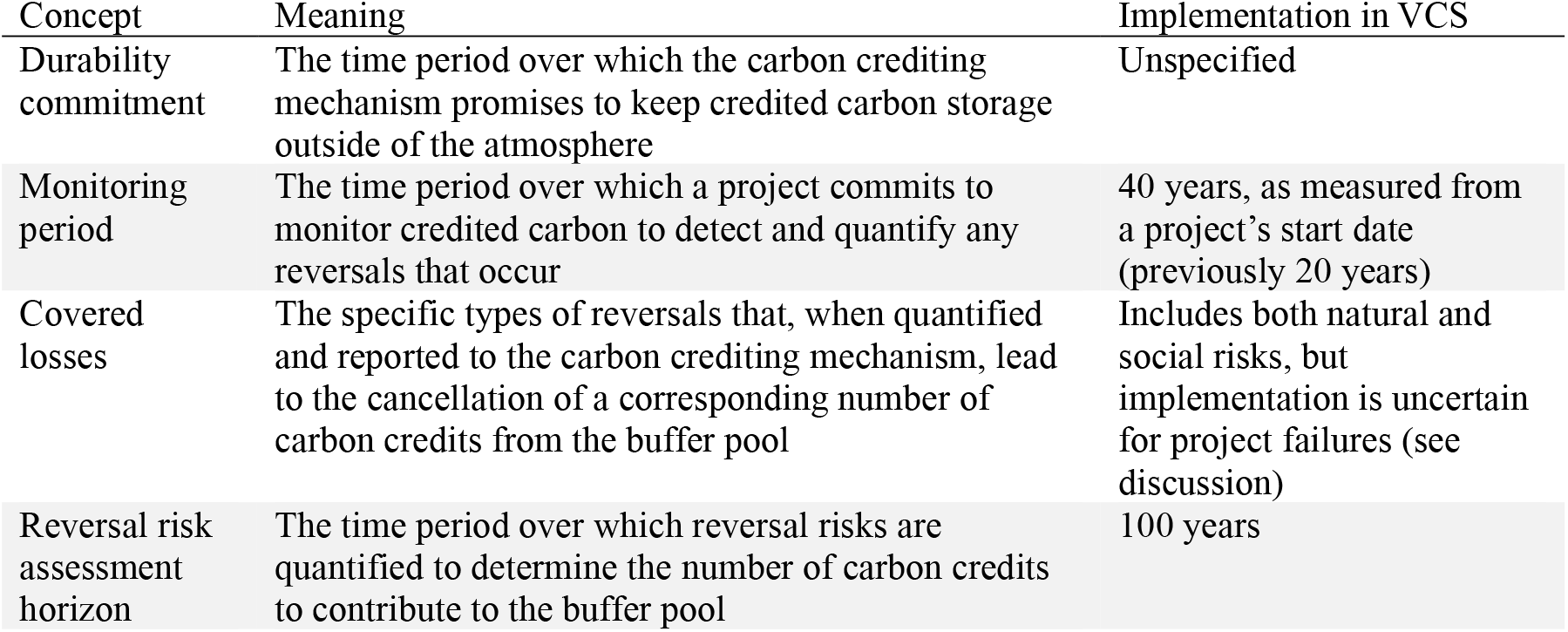
Concepts in reversal risk management.

| Concept | Meaning | Implementation in VCS |
| --- | --- | --- |
| Durability commitment | The time period over which the carbon crediting mechanism promises to keep credited carbon storage outside of the atmosphere | Unspecified |
| Monitoring period | The time period over which a project commits to monitor credited carbon to detect and quantify any reversals that occur | 40 years, as measured from a project’s start date (previously 20 years) |
| Covered losses | The specific types of reversals that, when quantified and reported to the carbon crediting mechanism, lead to the cancellation of a corresponding number of carbon credits from the buffer pool | Includes both natural and social risks, but implementation is uncertain for project failures (see discussion) |
| Reversal risk assessment horizon | The time period over which reversal risks are quantified to determine the number of carbon credits to contribute to the buffer pool | 100 years |

Verra’s standardized reversal risk assessment tool (Verra, 2024a) separates natural risks (e.g., fire, insects, drought) from what we call social risks — external factors like political instability and land tenure insecurity, and internal factors like financial viability and enforcement against illegal logging. For each risk factor, projects assess both the risk itself and any mitigating actions taken (e.g., forest thinning to reduce fire risk; Levine et al., 2026).

Although the tool’s flexibility allows projects to estimate risks as low as 0%, Verra imposes a minimum total (natural plus social) risk of 12% (Anderegg et al. 2025). As a result, the buffer pool only covers reversals up to the greater of a project’s self-assessed risk or 12%. If actual reversal risks exceed this threshold, then the buffer pool may become insolvent, threatening Verra’s ability to substantiate the durability of its VCUs.

## Results

### Social Risks to Forest Carbon Offset Projects

To characterize social risks to forest carbon offset projects in Verra’s VCS registry, we developed a dataset of all ‘agriculture, forestry, and other land use’ (AFOLU) projects registered in VCS as of February 17^th^, 2026. We further limited our analysis to projects labelled ‘afforestation, reforestation, and restoration’ (ARR), ‘improved forest management’ (IFM), and ‘reduced emissions from deforestation and degradation’ (REDD), to exclude non-forest projects (Fig. 1A-B). In total, we identified 244 qualifying projects (Fig. 1A).

**Figure 1.**
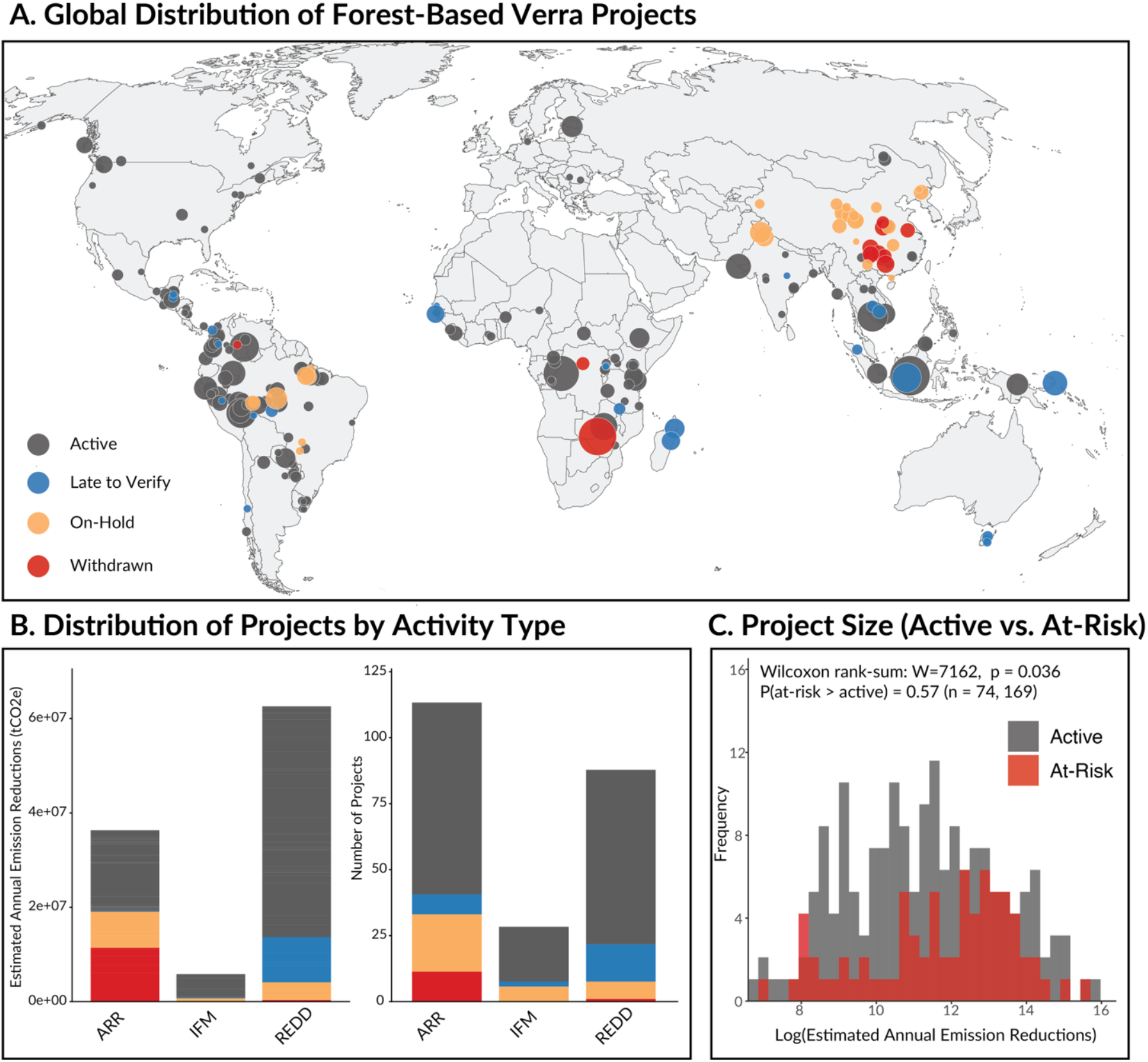
**A** Global distribution of forest-based carbon offset projects registered via Verra’s Voluntary Carbon Standard (VCS) as of February 17^th^, 2026. Projects are colored according to their status in Verra’s registry, where gray dots represent projects in good standing (‘active’), and colored dots represent ‘at-risk’ projects. **B** Distribution of VCS projects by activity type (ARR – afforestation, reforestation, and restoration; IFM – improved forest management; and REDD – reduced emissions from deforestation and degradation) and project status. The left panel characterizes this distribution by estimated annual emissions reductions (an indicator of project size), and the right panel by the number of projects in each category. **C** The distribution of active vs. at-risk project sizes (the logarithm of estimated annual emissions reductions). At-risk projects tend to be larger than active projects (Wilcoxon rank-sum test p-value = 0.036).

We then classified projects as ‘active’ (in good standing) or ‘at-risk’ (out of compliance) based on Verra’s internal status designations. At-risk statuses include: ‘withdrawn’, meaning the project no longer participates in the VCS program, either voluntarily or because Verra deemed it untrustworthy; ‘on-hold’, meaning Verra has identified a credible issue warranting further investigation; and ‘late-to-verify’, meaning the project missed the third-party verification VCS requires every five years to confirm net carbon benefits before issuing credits.

We define failure as a project ceasing its monitoring obligations before the end of its monitoring period, at which point the carbon storage represented by previously issued credits goes unmonitored and cannot definitively be relied upon. While not every at-risk designation results in failure, many at-risk projects are at high risk of or have already failed, so we use these labels as a systematic and transparent basis for risk assessment.

Of the 244 forest-based projects identified, 74 (30%) were at-risk, representing 22% of the total issued VCUs (Fig. 1). ARR projects had the highest proportion of projects at risk (36%), followed by IFM (27%) and REDD (25%), although IFM projects were relatively few (Fig. 1B). At-risk projects were larger on average than active projects, with a mean difference in estimated annual emissions reductions of 37,423 tCO_2_e year^-1^. These size distributions were statistically distinguishable (Wilcoxon rank-sum p = 0.036), and a randomly selected at-risk project had an approximately 57% probability of exceeding a randomly selected active project in size (Fig. 1C). Mean self-assessed social risk was 7.8%, and at-risk projects estimated higher social risks on average than active ones (9.7% vs. 7.1%; estimated mean difference = 2.58, bootstrapped CI = [0.482, 4.763]; Wilcoxon rank-sum test *p* = 0.0069; Figure S4).

### Time-dependent survival

To assess whether projects’ self-assessed social risks could cover realized rates of at-risk designation or failure, we conducted a time-dependent survival analysis. Because projects rarely receive an at-risk designation immediately—there is typically a delay of at least five years, corresponding to the first verification report’s due date—simply calculating the current share of at-risk projects would understate true social risk. Modeling the probability of transitioning into at-risk status as a function of project age instead yields unbiased estimates over defined time horizons.

At-risk status, however, is not equivalent to permanent project failure. Some projects return to good standing, for example if investigation of an on-hold project finds no wrongdoing, or delayed verification confirms no reversal occurred. We therefore model failure as a two-stage process: the age-dependent probability of entering at-risk status, followed by the probability that an at-risk project ultimately fails. Since the second transition cannot yet be precisely quantified with available data, we scale projected at-risk exposure by assumed failure rates of 25%, 50%, and 75% to bound plausible scenarios. We later analyze a limited longitudinal dataset to understand which scenario best matches observed outcomes.

The survival analysis indicates that projects’ self-assessed social risks are inadequate to address reversal risks over a 100-year time horizon. Even under the most optimistic scenario, the probability of a given project failing within its first 25 years (Fig. 2; Table 2) is nearly double the mean social risk estimated by individual projects (15.5% vs. 7.8%) over a period four times longer, and 1.5 times the mean total risk attributable to social factors (10.1%) after accounting for the 12% minimum allowable contribution. Overall, over half of projects (61.8%; Table 2) are expected to receive an at-risk designation within 25 years of the start of their crediting period.

**Table 2:** Projected failure rates and buffer pool impacts

| *Failure rate scenario* | *25-year cumulative failure probability* | *Projected share of projects failed by 2051* | *Projected buffer pool liability (% of current capitalization)* |
| --- | --- | --- | --- |
| Optimistic (25%) | 15.5% [10.0%, 18.9%] | 19.5% [14.4%, 24.6%] | 124% [82%, 166%] |
| Intermediate (50%) | 31.0% [20.2%, 37.8%] | 39.1% [28.8%, 49.3%] | 249% [165%, 332%] |
| Pessimistic (75%) | 46.3% [30.3%, 56.6%] | 58.6% [43.2%, 73.9%] | 373% [247%, 499%] |

**Figure 2.**
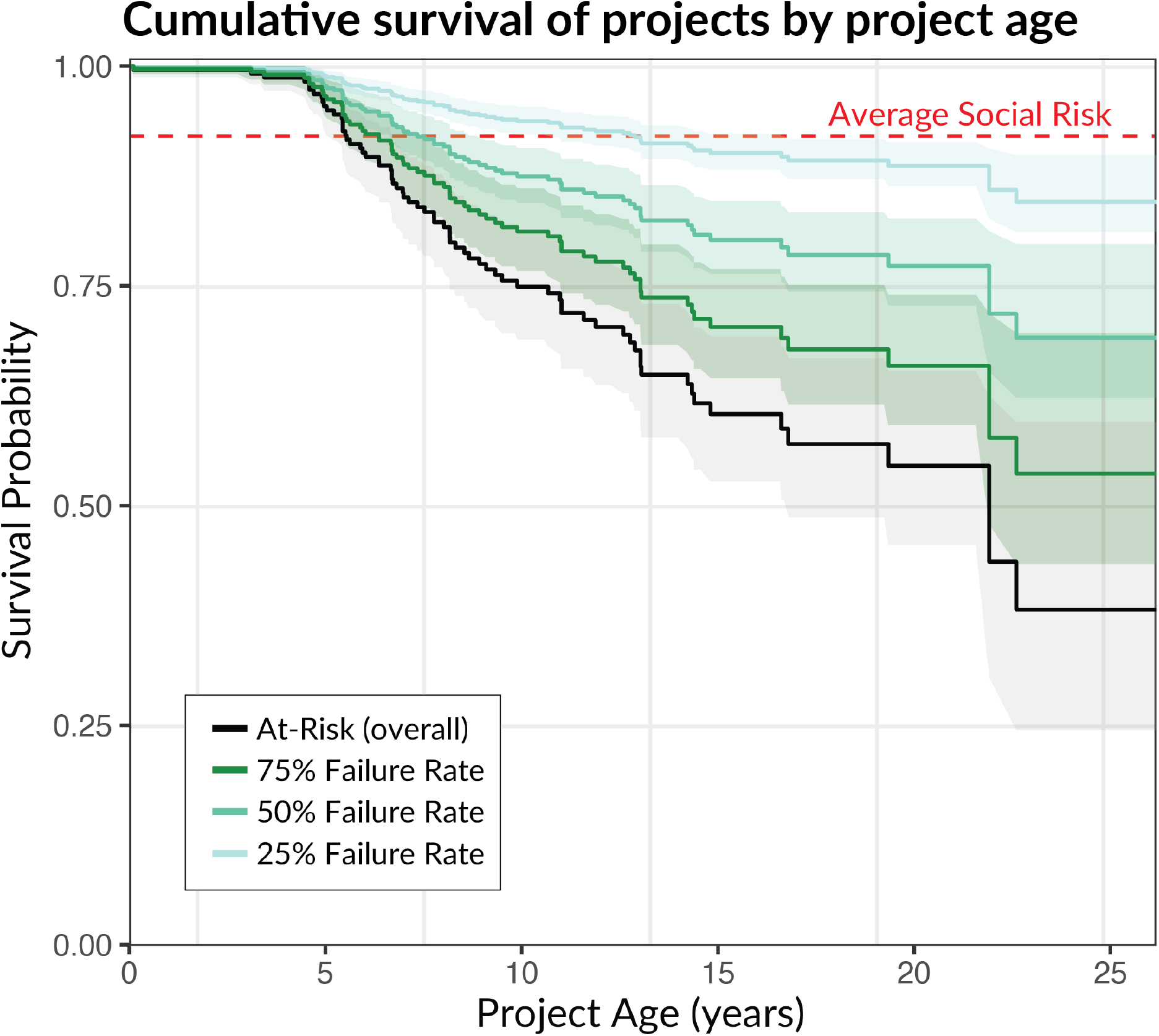
Cumulative survival of VCS projects through time. The black line shows the trajectory of at-risk designations through time for projects currently designated at-risk, with the shaded portion representing the 95% confidence interval. The colored lines represent estimated project failures when applying three different hypothetical failure rates: 25%, 50%, and 75%. The dashed red line indicates the mean 100-year survival rate estimated by individual projects in the VCS registry (both at-risk and active).

### Projecting future project failures and buffer pool impacts

Next, we used these survival estimates to forecast the accumulation of project failures through February 2051 (Methods; Fig. S2), comparing the total credits issued as of February 2026 to projects expected to fail, with the buffer pool capitalization as of the same date. Although both the buffer pool and number of issued credits will expand over time, this comparison is valid as the present-day buffer pool is intended to ensure the durability of all previously issued credits, while future credits will be covered by future contributions (Badgley et al., 2022). Across all three plausible failure rate scenarios, projected buffer pool liability exceeds current capitalization by 2051 (Fig. 3) and substantially outpaces the 75% of the pool dedicated to social risk, leaving no room for natural reversals (Table 2).

**Figure 3.**
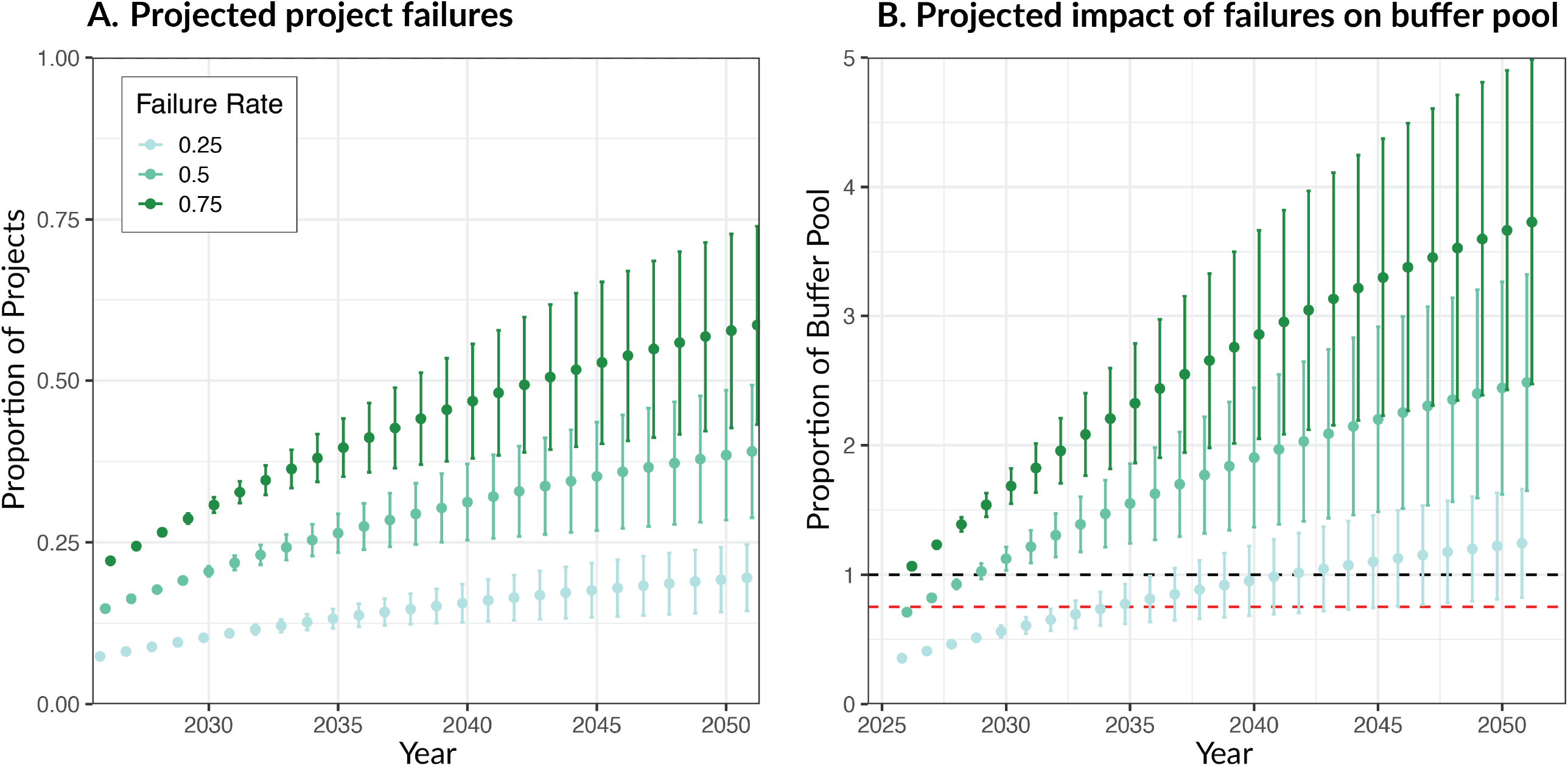
Projected project failures and associated buffer pool impacts. Panel A shows the projected proportion of projects expected to fail over time under three hypothetical failure realization rates (25%, 50%, and 75%), with points indicating mean estimates and error bars representing 95% confidence intervals. Panel B shows the corresponding projected impact of these failures on the buffer pool, expressed as the proportion of total buffer contributions at risk. The dashed black line indicates full buffer exposure (i.e., complete loss), while the dashed red line represents the contribution attributable to social risks alone.

### Assessing project failure rates with longitudinal data

To assess which failure rate scenario best matches realized outcomes in VCS, we used a longitudinal dataset of project status changes scraped from the VCS registry between February 2024 and February 2026 (G. Badgley, personal communication, 2026). For each at-risk project during this period, we identified its date of first at-risk designation and tracked subsequent status changes to one of three terminal outcomes: withdrawal (a proxy for failure), recovery (return to active status), or censoring (still at-risk as of February 2026). We then estimated the probability of withdrawal versus recovery over time using a competing risks analysis, which accounts for projects that had not yet reached either outcome by the end of the observation period.

Of the 22 projects (out of 83 total) that changed status, 57% recovered while the remaining 43% withdrew (Fig. S3). However, many recoveries belonged to a group of related projects placed on hold for minor reporting omissions rather than formal Verra investigations, unlike the on-hold projects in our primary dataset, which all arose from formal investigations. Excluding this group, only 32% recovered, versus 68% that withdrew (Fig. S3). Assuming recovery probability does not change substantially beyond two years of at-risk status, these results are consistent with the 50% and 75% failure rate scenarios and suggest failure rates below 25% are unlikely. If anything, this inference likely underestimates true failures, since our scenarios were applied even to withdrawn projects, which typically represent realized failures. While withdrawal doesn’t preclude re-registration outside VCS, which would relieve the VCS buffer pool of liability for that project’s credits, we found no evidence of re-registration among the 13 withdrawn projects in our dataset.

### Bounding the social-risk contribution required for durability

Using our estimates of at-risk designation rates (Fig. 2), we performed a preliminary extrapolation to project that sustaining currently observed at-risk-to-failure rates (0.43-0.68) without exhausting the VCS buffer pool would require a social-risk contribution of roughly 34-68% of issued credits—three to seven times the current average (10.1%). We caution that because these calculations require extrapolating fitted hazard models well beyond our 25-year dataset (Methods), they are intended to be illustrative rather than a precise guide for policy.

Nonetheless, these numbers are consistent across hazard models and time horizons (40 and 100 years; Table S2), suggesting that the gap is unlikely to close with incremental adjustments to current minimums.

## Discussion

Our analysis of Verra’s forest-based carbon offset portfolio suggests the existing buffer pool is insufficient to cover social risks under a range of plausible failure scenarios based on 2000-2026 performance. Even under optimistic failure rates, the volume of credits issued to failed projects exceeds the buffer pool’s total size by the early 2040s and greatly exceeds the portion allocated to social risks (Fig. 3; Table 2). This echoes findings of substantial reversal vulnerability across other markets, registries, and regions (Anderegg et al., 2025; Badgley et al., 2022; Galik & Jackson, 2009; Haya et al., 2023; Wu et al., 2023, 2026).

A key uncertainty, however, is that the implications of project failure for the buffer pool depend on how its purpose is defined. We assume the buffer pool is intended to ensure that issued credits correspond to real, maintained carbon storage. Under this interpretation, if a project fails, credits equivalent to its total issuance should be cancelled from the buffer pool, as the credited carbon would no longer be actively protected and thus the climate mitigation claim no longer guaranteed (Cabiyo & Field, 2025). This definition aligns with the Paris Agreement Crediting Mechanism, which automatically deems project failures to be reversals (UNFCCC, 2025a).

### Changes in VCS program rules and operations

Earlier VCS versions treated project failures as reversals. Under VCS Standard v4.5 (valid through February 4^th^, 2026), failure to submit verification triggered cancellation of credits equal to total issuance after a 15-year waiting period (Verra, 2024b). The current VCS Standard (v5.0) removed this requirement, implicitly on the theory that failure does not necessarily lead to a physical reversal. Instead, after a shorter 10-year waiting period, only the projects’ buffer contribution—approximately 13.3% of issued credits on average—is cancelled (Verra, 2025b). Further cancellation only occurs if Verra detects a reversal using its new remote-sensing-based ‘long-term monitoring system,’ whose methodology and performance remain unclear. In effect, these changes shrink buffer pool cancellations; make further cancellation contingent on future reversal detection; and shift monitoring to a centralized Verra-run system, raising moral hazard concerns.

### A reluctance to draw on the buffer pool

This policy shift coincides with several high-profile VCS project failures. The most notable is the Kariba REDD+ project, the second largest in our database by estimated annual emissions reductions (and largest at-risk project; Table S1). In 2023, independent investigations revealed that Kariba had substantially overstated baseline deforestation rates, resulting in excess credit issuance (Blake, 2023; Elgin, 2025; Verra, 2025c). Following media scrutiny and a subsequent Verra investigation, the project was placed on hold and later withdrawn by its developer. Verra ultimately determined that the project had been issued approximately 15 million excess VCUs (Verra, 2025c), although it did not explain the basis for this calculation (Elgin, 2025). Rather than cancel Kariba’s issuance from its buffer pool, Verra asked the developer to replace them and requested that buyers voluntarily cancel remaining units. However, since the project exited the VCS program, it remains unclear whether the developer will comply without legal action, which has not yet occurred to our knowledge. If the excess credits aren’t replaced or cancelled from the buffer pool, buyers are left holding credits with no corresponding climate mitigation outcome (Badgley et al., 2025).

The Suruí Forest Carbon Project in Brazil illustrates a different form of failure. It has not been verified since 2013, likely related to mining-driven deforestation in 2014 and produced estimated emissions exceeding the project’s total credit issuance (Verra, 2018). Rather than quantifying this reversal as required, Suruí has simply ceased participation in VCS without formally withdrawing. VCS rules do not clearly address this scenario, but a strict reading suggests only the project’s buffer contribution would be cancelled, leaving most of the reversal unaddressed and the atmosphere on the hook—unless Verra were to retroactively detect the reversal via its promised ‘long-term monitoring system’ (Verra, 2025b). Similarly, Verra recently cancelled four China-based projects for lacking government authorization and placed 35 related projects on hold. As with Kariba, Verra appears to be pursuing developer remediation rather than buffer cancellation (Verra, 2025a).

Together, these case studies and VCS rule changes suggest Verra may be hesitant to use its buffer pool to address project failures arising from social risks and is instead relying on voluntary credit cancellations and VCU replacement by developers. As a result, credit durability rests on developers’ willingness to repurchase millions of dollars in third-party VCUs, or on Verra’s ability to legally enforce its requirements. We are not aware of either approach succeeding to date. However, our finding that the entire buffer pool could be threatened within fifteen years should be interpreted cautiously: if Verra successfully extracts compensation, the cancellations we project may not materialize; but if Verra is unsuccessful, the buffer pool is unlikely to withstand the forecast failure rate.

### Policy recommendations and conclusion

Our findings suggest three changes to improve the VCM’s resilience to social risks. First, the minimum buffer pool contribution for social risks should be increased. Even under the most optimistic failure rate assumption, 15% of projects are projected to fail within 25 years due to social factors (Fig. 2). This exceeds both the average self-assessed social risk (7.8%) and the average overall buffer contribution (13.3%). Our illustrative projections indicate that contributions may need to increase as much as three- to seven-fold to address 100 years of social risks. Second, rather than relying on project developers to replace VCUs in the case of social failure, registries should treat project failures as reversals, cancel buffer credits equivalent to total issuance, and then seek compensation from project developers. Doing so ensures credit validity while the uncertain, potentially years-long repayment process unfolds and follows precedent under the Paris Agreement Crediting Mechanism (UNFCCC, 2025a, 2025b). While cancelling credits equal to total issuance could be conservative, this approach may be preferable to satellite-based detection, which is difficult in dense, mature forests (Gao et al., 2020; Xu et al., 2025).

Third, more attention should be given to strategies that reduce the risk of uncompensated project failures, such as increasing the administrative capacity to enforce VCM replacement rules, implementing risk-sharing measures such as bonding requirements or third-party guarantees, treating forest credits as temporary or mitigation contribution credits as under the UN Clean Development Mechanism, or requiring financially independent verification bodies to carefully assess social risks.

## Materials and Methods

### VCS data

We assembled a dataset of all forest-based VCS projects by querying Verra’s online registry (registry.verra.org) for projects labelled ‘agriculture, forestry, and other land use’ (AFOLU) on February 17^th^ 2026. We additionally filtered these projects for those with activities labelled ‘afforestation, reforestation, and restoration’ (ARR), ‘improved forest management’ (IFM), and ‘reduced emissions from deforestation and degradation’ (REDD), to ensure our analysis was limited to forest-based projects. We also pulled the records of all VCU issuances and buffer contributions for AFOLU projects and matched these to the project data. Finally, we used these data to exclude projects which have not yet been issued credits.

To determine each projects’ sequence of verifications we pulled all verification reports available for projects in our dataset and recorded the most recent date of issue for each verification report. While Verra bases verification deadlines on the date of approval, which often occurs several months after the date of issue, approval dates are not publicly available and so we used date of issue as a proxy to establish project timelines. We then pulled data on each project’s reversal risk assessment from the non-permanence risk reports published on each project’s page in the online VCS registry. We separately recorded internal, external, and natural risks, as well as each project’s effective overall risk score which was equal to either the sum of internal, external, and natural risks, rounded to the nearest whole number, or twelve if the sum of risks was less than this minimum. Finally, we established the date at which each at-risk project (projects with statuses of ‘late-to-verify,’ ‘withdrawn,’ or ‘on-hold’) gained at-risk status. For late-to-verify projects, we simply used five years after the most recent verification date. For on-hold projects, we used the date listed in the ‘new review notification letter,’ which is posted on the project’s registry page following a change in status to on-hold. For ‘withdrawn’ projects, which have no standardized source of information on withdrawal date, we searched for reports of the withdrawal, either from Verra itself or independent news sources.

### Project characteristics

To quantify differences in project size and self-assessed risk between at-risk and active projects we compared both the means and distributions of these variables. We used each project’s estimated annual emissions reductions as a proxy for project size, as this variable is closely tied to the number of credits the project is eligible to receive. For self-assessed risk, we used the sum of projects’ internal and external risks reported in the non-permanence risk report. To compare mean project size and social risk between at-risk and active projects, we used a nonparametric bootstrap with 2,000 iterations to construct a confidence interval for the mean difference in each metric. Differences in project size and social risk were further evaluated using a one-sided Wilcoxon rank-sum test, a nonparametric alternative to the two-sample t-test. Effect size was quantified using the ‘probability of superiority,’ defined as the probability that a randomly selected at-risk project has a larger size (e.g. higher social risk) than a randomly selected active project. All statistical tests were conducted in R (version 4.4.1; R Core Team, 2013).

### Survival analysis

We conducted a survival analysis to assess the observed risk of social failure among projects in the VCS database. First, we constructed a time-to-event dataset using each projects’ at-risk and crediting period start dates. For each project, the observation period began at the start of the crediting period and ended either at the date the project transitioned to at-risk status or at a fixed censoring date (February 17^th^, 2026; for active projects). Using the crediting period start date as a proxy for the true start of the observation period is imperfect, since projects frequently credit emissions reductions and removals retroactively. However, publicly available registration dates were unreliable and often contradicted other project documentation, making the crediting period start date the more consistent choice. However, we note that because crediting periods frequently begin before a project’s true start date, this choice overstates each project’s true exposure time, potentially causing us to underestimate social risks through time.

The survival analysis was conducted using the Kaplan-Meier estimator to assess the probability of remaining active over time (Goel et al., 2010). Cumulative at-risk probabilities were derived as one minus the survival function. To account for uncertainty in the rate at which at-risk projects ultimately fail, we considered three possible failure rates for at-risk projects: 25%, 50%, and 75%. Adjusted survival curves were obtained by proportionally scaling cumulative at-risk probabilities, with confidence intervals from the Kaplan-Meier estimator propagated through these transformations. Finally, we compared these adjusted survival estimates to the average self-assessed social risk of projects in our database to determine whether project’s self-assessments aligned with realized risks. The survival analysis was conducted using the ‘survival’ package in R (Therneau, 2026).

In addition to the standard survival analysis, we also performed a competing risks analysis to determine the relative risks of transitioning to each of the at-risk statuses: late-to-verify, withdrawn, and on-hold. We further subdivided the late-to-verify status according to whether projects were more or less than two years late to submit verification (i.e., seven years since most recent verification). We did so as projects which are more than two years late are less likely to return to active status. Active projects were again treated as right-censored observations at the administrative censoring date (February 17^th^, 2026). We estimated cause-specific cumulative incidence functions using nonparametric methods as implemented using the ‘cmprsk’ package in R (Bob Gray, 2025), which account for the presence of competing events. We then used variance estimates from the cumulative incidence estimator were to construct approximate confidence intervals (Fig. S1).

### Projecting future project failures

To extrapolate project failures into the future, we fit a series of parametric survival models to the dataset described above, including the following functional forms: Weibull, lognormal, Gompertz, and a flexible spline-based model. All models were fit using the ‘flexsurv’ package (Jackson, 2016). Model fit was compared using the Akaike Information Criterion and by visually comparing fitted survival curves against the nonparametric Kaplan-Meier estimate (Fig. S2). The best-fitting model, the flexible spline, was used for forward projections.

Projected failures among currently active projects were estimated by computing, for each project, the conditional probability of failure over each of the next 25 years given survival to its observed age. Expected failures at each horizon were then obtained by summing these conditional probabilities across active projects and adding projects already classified as at-risk. To reflect uncertainty about the fraction of at-risk projects that will ultimately fail, we again scaled projected totals by three failure rates (25%, 50%, and 75%). Confidence intervals for projected failures were obtained using the delta method, propagating uncertainty in the spline model’s parameter estimates (Oehlert, 1992).

The same method was used to estimate the proportion of the buffer pool threatened by project failures through 2051. For each active project, the expected future buffer loss was calculated as the product of its conditional failure probability (scaled by the three failure rates) and its current number of issued credits. These project-level expected losses were then summed across projects. Existing at-risk projects were incorporated directly into these calculations by scaling them according to each of the three failure rate scenarios. Uncertainty was again quantified using confidence intervals derived from the delta-method.

### Longitudinal analysis of at-risk project outcomes

To characterize project trajectories following entry into at-risk status, we relied on a longitudinal dataset of project status in the VCS registry (G. Badgley, personal communication, 2026), which was constructed by regularly scraped registry data, beginning in February 2024. We then combined these snapshots with archived records to reconstruct project-level status histories. Where necessary, at-risk entry dates were supplemented using documentation from the VCS registry (e.g., monitoring reports and notification letters).

Projects were included if they were classified as at risk (e.g., “late to verify,” “on hold,” or “withdrawn”) at any point between February 2024 and February 2026. For each project, we defined the at-risk entry date as the earliest documented transition into an at-risk state using the same methodology described above for the primary dataset. We then classified three terminal outcomes: (1) failure, defined as withdrawal from the VCS registry; (2) recovery, defined as return to good standing (e.g., “registered” or “active”); and (3) censoring, defined as projects remaining at risk at the end of the observation period. Outcome dates were defined as the first observed transition to each state.

Time-to-event was measured from at-risk entry to failure, recovery, or censoring. We estimated the cumulative incidence of failure and recovery using competing risks methods implemented in the cmprsk package in R (Bob Gray, 2025).

Because a subset of recovering projects belonged to a related group placed on hold due to minor reporting omissions rather than formal Verra investigations, we conducted a sensitivity analysis excluding these projects. Results were compared across specifications to assess robustness and to evaluate consistency with assumed failure rate scenarios used in the main analysis.

### Estimating minimum social risk contributions required for durability

To estimate the social-risk buffer pool contribution required to ensure solvency under currently observed rates of at-risk designation and project failure rates, we inverted our survival model estimates to solve for the social-risk contribution required to ensure the buffer pool is robust to social reversals.

We first defined the buffer pool’s social-risk budget as 10.1%, which is the share of the buffer pool attributable to social risks. To calculate this number, we summed internal and external risks across all projects and divided by the sum of all assessed risks (internal, external, and natural), giving the portfolio-wide proportion of assessed risk attributable to social factors (75.6%). We then multiplied this proportion by the mean effective risk score across projects (13.3%), which is the mean buffer pool contribution projects were required to make. We note that projects assessed under earlier versions of Verra’s risk tool were subject to a 10% minimum rather than the current 12%, so the mean effective risk score reflects the minimum in force at the time of each assessment; it closely matches the buffer pool’s realized capitalization of 13.2% of gross issuance. Next, we used each of our four fitted survival models (Weibull, lognormal, Gompertz, and spline; see above) to estimate the cumulative probability that a project becomes at-risk by 40 years, and 100 years. These two horizons were chosen because they reflect VCS’s monitoring period length and reversal risk assessment horizon, respectively (Table 1).

We then solved for the social-risk budget required to ensure durability under each time horizon given our empirically observed at-risk-to-failure rate. For a given horizon, the cumulative expected failure of VCS projects is the product of the probability that a project becomes at-risk by that date, and the probability that an at-risk project ultimately fails. To quantify the second of these terms, the at-risk-to-failure rate, we rely on the strong assumption that a single rate applies uniformly across projects and project ages. We then used our empirically observed at-risk-to-failure rates from the longitudinal analysis (43% including, and 68% excluding the group of related projects placed on hold for minor reporting issues) and report the resulting required budget as a both a raw percentage value and multiple of the current social risk budget.

Because both the 40- and 100-year time horizons lie well beyond the reach of our dataset (approximately 25 years), estimating the rates of at-risk designation at these times requires extrapolating each fitted hazard model well past the range of observed data. This extrapolation is a major source of instability in survival analysis, as different functional forms which fit the observed data similarly can diverge substantially beyond this range, particularly for forms with differing asymptotic behavior. For example, the Weibull (with shape parameter greater than one) and Gompertz hazard functions increase without bound, and both models estimated a probability of at-risk designation close to 1 for both horizons. However, because the spline model, which is capable of representing a saturating hazard trajectory, produced similar estimates, the estimates of near-100% hazard does not appear to be solely the result of assuming monotonically increasing hazard functions. However, the spline model’s extrapolated behavior may itself be unreliable, a general limitation of flexible hazard models extended beyond their fitted range. Therefore, the convergence of results across models at long time horizons is not a reliable indicator of accuracy. For these reasons, we used multiple hazard functions and view this extrapolation as illustrative, rather than a quantitative forecast.

Another key source of uncertainty comes from assuming that the at-risk-to-failure rates observed in our 2-year longitudinal dataset will remain constant over decades and across projects. This is unlikely to be true. Therefore, we report our estimates of minimum social risk contributions as an illustrative range across models and failure rates rather than as a single point estimate. We strongly caution against including these numbers into policy directly.

## Acknowledgments

We are grateful to Grayson Badgley for sharing his longitudinal dataset of VCS project statuses, as well as for his feedback on our analysis and approach. Data collection was supported in part by Lincoln Bell, Adam Kergaye, and Felix Pollock. We also thank the members of the Anderegg lab at the University of Utah for their helpful feedback on earlier versions of the manuscript. JIL was supported by the Wilkes Postdoctoral Fellowship awarded by the Wilkes Center for Climate Science and Policy at the University of Utah. WRLA acknowledges support from the David and Lucille Packard Foundation, US National Science Foundation grants 2003017, 2044937, 2330582, 2519523 as well as the Alan T. Waterman award IOS-2325700. DC is a member of the Paris Agreement Crediting Mechanism’s Methodological Expert Panel, but does not speak for the panel or the mechanism here.

## Notes

### Competing Interest Statement

The authors have declared no competing interest.

## References

Allen, M. R., Frame, D. J., Friedlingstein, P., Gillett, N. P., Grassi, G., Gregory, J. M., Hare, W., House, J., Huntingford, C., Jenkins, S., Jones, C. D., Knutti, R., Lowe, J. A., Matthews, H. D., Meinshausen, M., Meinshausen, N., Peters, G. P., Plattner, G.-K., Raper, S., … Zickfeld, K. (2025). Geological Net Zero and the need for disaggregated accounting for carbon sinks. Nature, 638(8050), 343–350. 10.1038/s41586-024-08326-8

Allen, M. R., Friedlingstein, P., Girardin, C. A. J., Jenkins, S., Malhi, Y., Mitchell-Larson, E., Peters, G. P., & Rajamani, L. (2022). Net Zero: Science, Origins, and Implications. Annual Review of Environment and Resources, 47(Volume 47, 2022), 849–887. 10.1146/annurev-environ-112320-105050

Anderegg, W. R. L., Trugman, A. T., Vargas G. G., Wu, C., & Yang, L. (2025). Current Forest Carbon Offset Buffer Pool Contributions Do Not Adequately Insure Against Disturbance-Driven Carbon Losses. Global Change Biology, 31(6), e70251. 10.1111/gcb.70251

Badgley, G., Allen, C., & Chay, F. (2025). Verra doesn’t have a good solution for excess crediting. CarbonPlan. https://carbonplan.org/research/kariba-excess-credits

Badgley, G., Chay, F., Chegwidden, O. S., Hamman, J. J., Freeman, J., & Cullenward, D. (2022). California’s forest carbon offsets buffer pool is severely undercapitalized. Frontiers in Forests and Global Change, 5. 10.3389/ffgc.2022.930426

Blake, H. (2023, October 16). The Great Cash-for-Carbon Hustle. The New Yorker. https://www.newyorker.com/magazine/2023/10/23/the-great-cash-for-carbon-hustle

Bob Gray. (2025). cmprsk: Subdistribution Analysis of Competing Risks (p. 2. 2-12) [Dataset]. 10.32614/CRAN.package.cmprsk

Brunner, C., Hausfather, Z., & Knutti, R. (2024). Durability of carbon dioxide removal is critical for Paris climate goals. Communications Earth & Environment, 5(1), 645. 10.1038/s43247-024-01808-7

Cabiyo, B., & Field, C. B. (2025). Embracing imperfection: Carbon offset markets must learn to mitigate the risk of overcrediting. PNAS Nexus, 4(5), pgaf091. 10.1093/pnasnexus/pgaf091

Cullenward, D., Badgley, G., & Chay, F. (2023). Carbon offsets are incompatible with the Paris Agreement. One Earth, 6(9), 1085–1088. 10.1016/j.oneear.2023.08.014

Elgin, B. (2025, October 17). Majority of Carbon Credits From Tarnished Project Deemed Bogus. Bloomberg.Com. https://www.bloomberg.com/news/articles/2025-10-17/majority-of-carbon-credits-from-tarnished-project-deemed-bogus

Galik, C. S., & Jackson, R. B. (2009). Risks to forest carbon offset projects in a changing climate. Forest Ecology and Management, 257(11), 2209–2216. 10.1016/j.foreco.2009.03.017

Gao, Y., Skutsch, M., Paneque-Gálvez, J., & Ghilardi, A. (2020). Remote sensing of forest degradation: A review. Environmental Research Letters, 15(10), 103001. 10.1088/1748-9326/abaad7

Goel, M. K., Khanna, P., & Kishore, J. (2010). Understanding survival analysis: Kaplan-Meier estimate. International Journal of Ayurveda Research, 1(4), 274–278. 10.4103/0974-7788.76794

Haya, B. K., Anderegg, W. R. L., Beymer-Farris, B., Blanchard, L., & Bomfim, B. (2023). Quality Assessment of REDD+ Carbon Credit Projects. Berkeley Carbon Trading Project.

Jackson, C. (2016). flexsurv: A Platform for Parametric Survival Modeling in R. Journal of Statistical Software, 70, 1–33. 10.18637/jss.v070.i08

Kreibich, N. (2024). Toward global net zero: The voluntary carbon market on its quest to find its place in the post-Paris climate regime. WIREs Climate Change, 15(5), e892. 10.1002/wcc.892

Levine, J. I., Zahnd, C., Blanchard, L., Fickle, J. C., Lengyel, T., Liu, M., Post-Leon, A. C., Wu, C., Xu, H., Yang, L., & Anderegg, W. R. L. (2026). Variable impacts of forest treatments on carbon and mortality following disturbance. Environmental Research Letters, 21(5), 053002. 10.1088/1748-9326/ae4888

Marland, G., Fruit, K., & Sedjo, R. (2001). Accounting for sequestered carbon: The question of permanence. Environmental Science & Policy, 4(6), 259–268. 10.1016/S1462-9011(01)00038-7

Michaelowa, A., Keßler, J., Dalfiume, S., & Ahonen, H.-M. (2025). Reversal risk and buffer pool contribution analysis. Perspectives Climate Group GmbH. Retrieved from Https://Www.Carbon-Mechanisms.de/Fileadmin/Media/Dokumente/PCG_Reversal_Risk_Paper_20250616.Pdf.

Oehlert, G. W. (1992). A Note on the Delta Method. The American Statistician, 46(1), 27–29. 10.1080/00031305.1992.10475842

Probst, B. S., Toetzke, M., Kontoleon, A., Díaz Anadón, L., Minx, J. C., Haya, B. K., Schneider, L., Trotter, P. A., West, T. A. P., Gill-Wiehl, A., & Hoffmann, V. H. (2024). Systematic assessment of the achieved emission reductions of carbon crediting projects. Nature Communications, 15(1), 9562. 10.1038/s41467-024-53645-z

R Core Team, R. (2013). R: A language and environment for statistical computing.

Romm, J., Lezak, S., & Alshamsi, A. (2025). Are Carbon Offsets Fixable? Annual Review of Environment and Resources, 50(Volume 50, 2025), 649–680. 10.1146/annurev-environ-112823-064813

Ruiting, W., Ahmad, M. S., Bin Jamaludin, J., Rahim, N. A., Wong, S. Y., Rehman, J. U., Kumar, L., & El Haj Assad, M. (2025). A comprehensive bibliometric review of voluntary carbon markets: Trends and future directions. Environmental Research Communications, 7(4), 042001. 10.1088/2515-7620/adc659

Tang, Y., Yang, C., Wu, H., Xu, Z., Tan, L., Tu, W., Li, B., Li, Z., Wang, Z., Guo, K., Xiong, S., Chen, S., Zhang, B., Tian, J., Hu, Y., Chen, Z., Chase, J. M., & Li, Q. (2025). Tropical forest carbon offsets deliver partial gains amid persistent over-crediting. Science, 390(6769), 182–187. 10.1126/science.adw4094

Therneau, T. (2026). A package for survival analysis in R.

Trouwloon, D., Streck, C., Chagas, T., & Martinus, G. (2023). Understanding the Use of Carbon Credits by Companies: A Review of the Defining Elements of Corporate Climate Claims. Global Challenges, 7(4), 2200158. 10.1002/gch2.202200158

UNFCCC. (2025a). Information note: Elements related to non-permanence and reversals for inclusion in relevant regulatory documents. https://unfccc.int/sites/default/files/resource/A6.4-INFO-METH-002.pdf

UNFCCC. (2025b). Standard: Requirements for activities involving removals under the Article 6.4 mechanism. https://unfccc.int/sites/default/files/resource/A6.4-STAN-METH-002.pdf

Verra. (2018). Surui Forest Carbon Project Loss Event. Verra.

Verra. (2024a). AFOLU Non-Permanence Risk Tool v4.2. Verra.

Verra. (2024b). Registration and Issuance Process, v4.6. Verra. https://verra.org/wp-content/uploads/2024/10/Registration-and-Issuance-Process-v4.6-1.pdf#page=32.15

Verra. (2025a). Serious Allegations Prompt Verra to Reject China Projects and Launch Broader Reviews. Verra. https://verra.org/serious-allegations-prompt-verra-to-reject-china-projects-and-launch-broader-reviews/

Verra. (2025b). VCS Registration and Issuance Process, v5.0. Verra. https://verra.org/wp-content/uploads/2025/12/Registration-and-Issuance-Process-v5.0.pdf#page=30.09

Verra. (2025c). Verra Acts on Kariba Project: Cancels Excess Credits, Advances Independent Review. Verra. https://verra.org/verra-acts-on-kariba-project-cancels-excess-credits-advances-independent-review/

West, T. A. P., Wunder, S., Sills, E. O., Börner, J., Rifai, S. W., Neidermeier, A. N., Frey, G. P., & Kontoleon, A. (2023). Action needed to make carbon offsets from forest conservation work for climate change mitigation. Science, 381(6660), 873–877. 10.1126/science.ade3535

Wu, C., Badgley, G., Goulden, M. L., Randerson, J. T., Trugman, A. T., Wang, J. A., Yang, L., Acil, N., Cook-Patton, S. C., Cullenward, D., Davis, S. J., Williams, C. A., & Anderegg, W. R. L. (2026). Forest carbon protocols underestimate climate-driven carbon loss risks. Nature, 654(8117), 107–113. 10.1038/s41586-026-10571-y

Wu, C., Coffield, S. R., Goulden, M. L., Randerson, J. T., Trugman, A. T., & Anderegg, W. R. L. (2023). Uncertainty in US forest carbon storage potential due to climate risks. Nature Geoscience, 16(5), 422–429. 10.1038/s41561-023-01166-7

Xu, W., Cheng, Y., Luo, M., Mai, X., Wang, W., Zhang, W., & Wang, Y. (2025). Progress and Limitations in Forest Carbon Stock Estimation Using Remote Sensing Technologies: A Comprehensive Review. Forests, 16(3). 10.3390/f16030449

