## Supplemental_Information for "Large and persistent social risks to forest carbon projects"

This file contains:

Supplemental Figures S1-S4

Supplemental Tables S1-S2

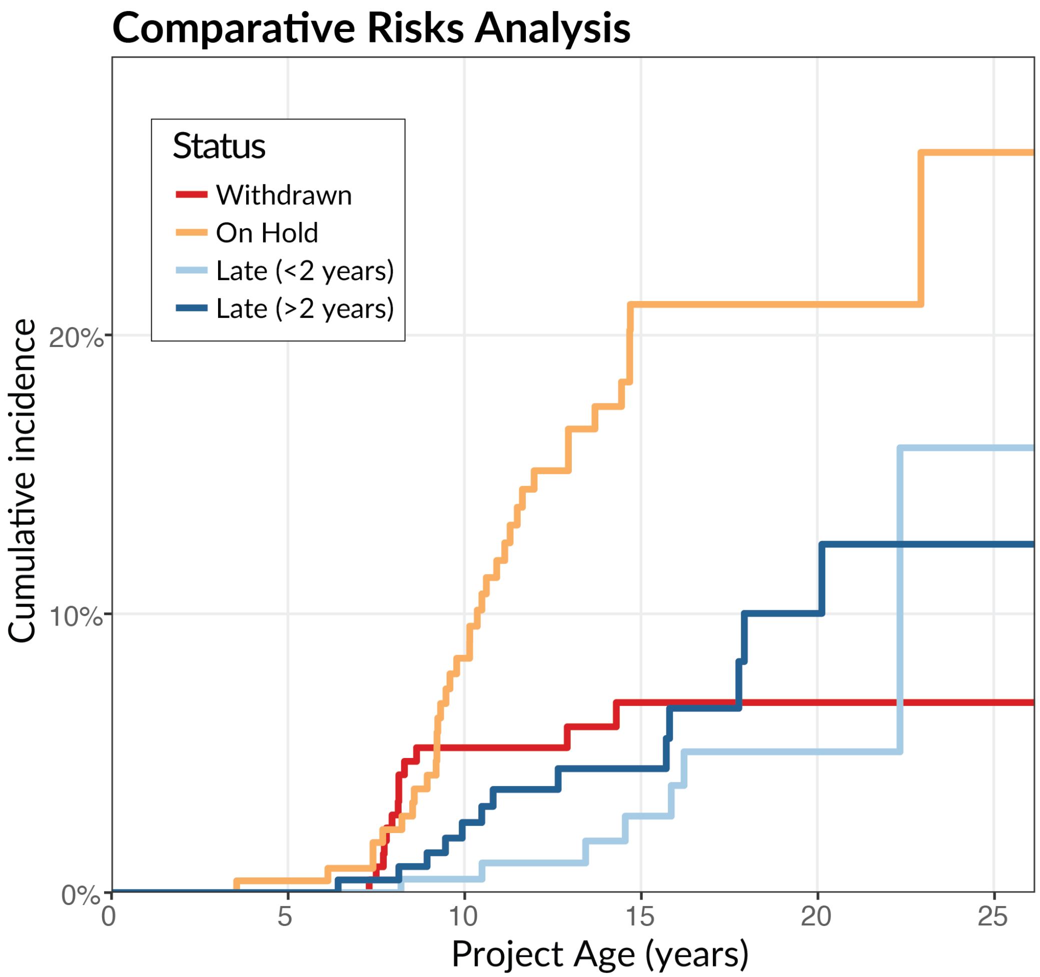


*Figure S1 – Cumulative incidence of project outcomes by type. The figure shows the cumulative incidence of distinct project outcomes—withdrawn, on hold, late (<2 years), and late (>2 years)—as a function of project age. Each step function represents the estimated probability that a project has transitioned into a given status over time, accounting for competing risks across outcome categories.*

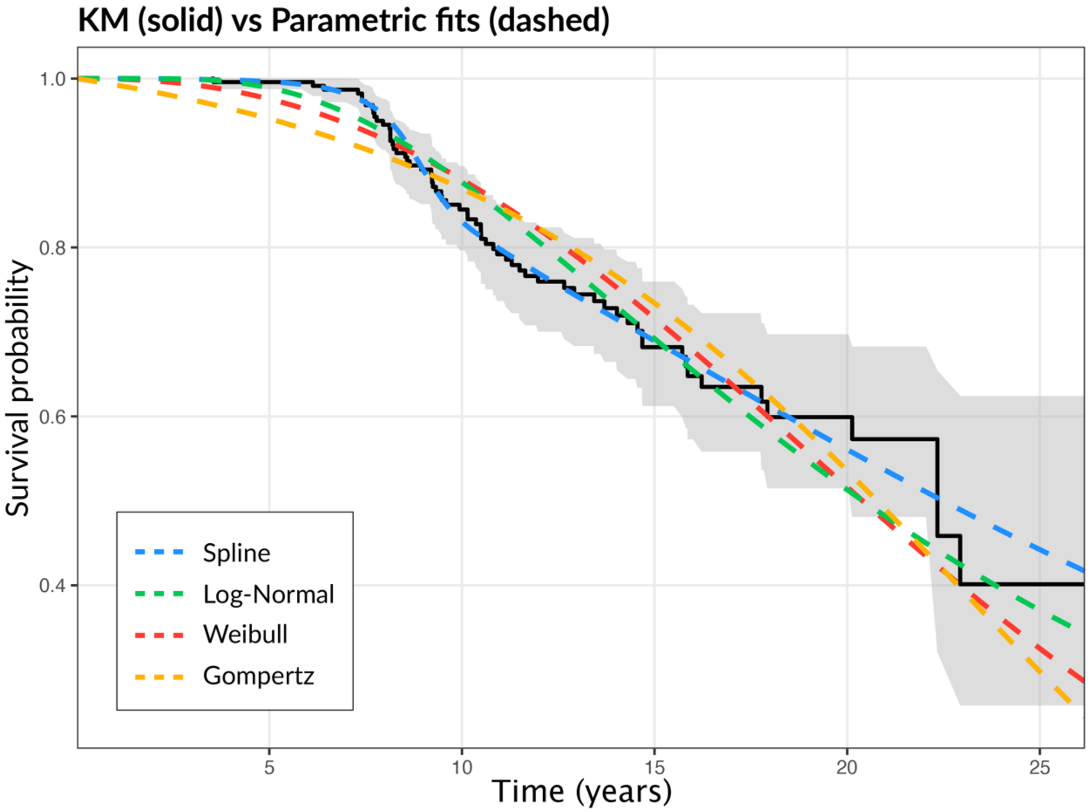


*Figure S2 – Kaplan–Meier survival estimates and parametric model fits. The solid black line shows the Kaplan–Meier estimate of project survival over time, with the shaded region indicating the 95% confidence interval. Dashed lines represent fitted parametric survival models, including spline-based, lognormal, Weibull, and Gompertz specifications.*

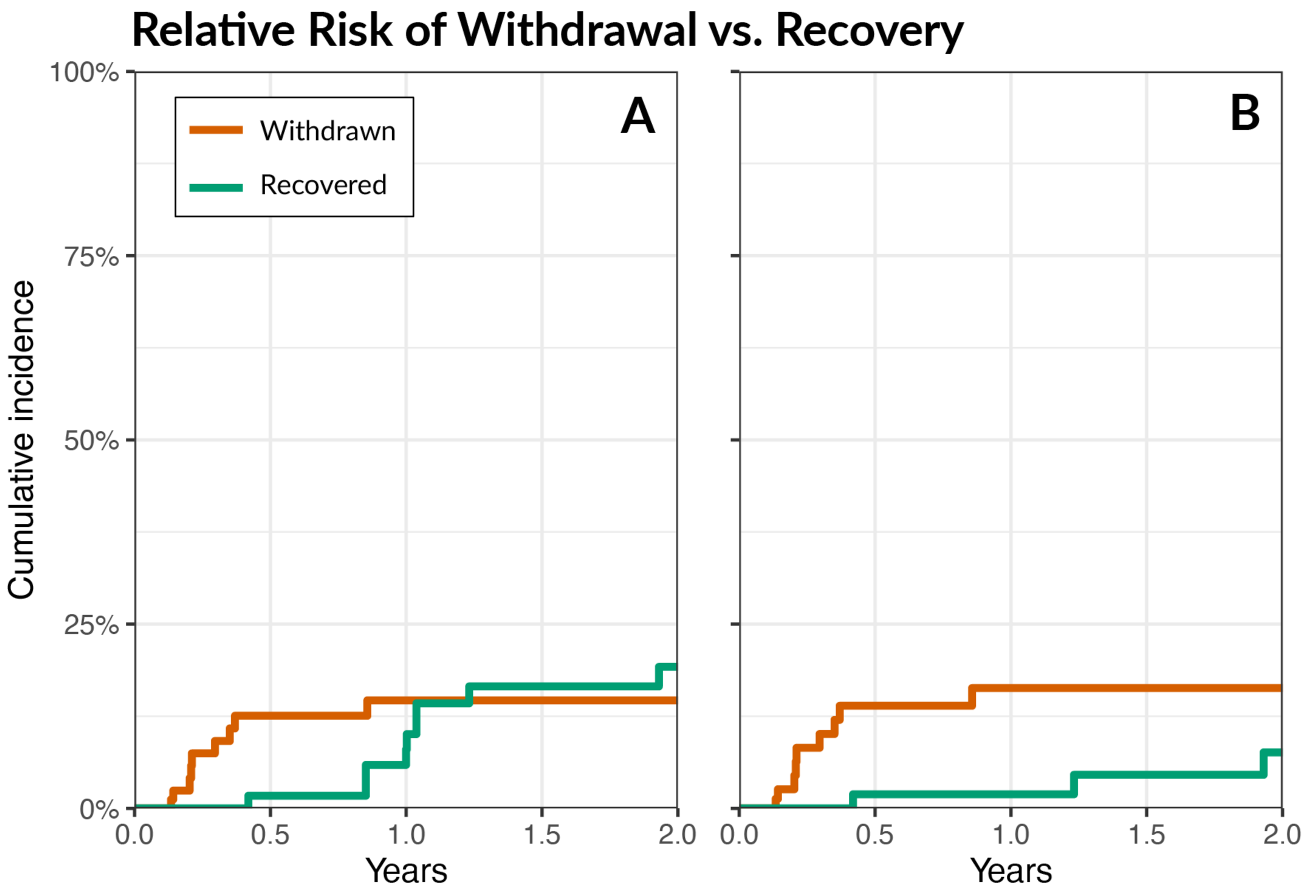


*Figure S3 -- Relative risk of project withdrawal vs. recovery from at-risk status. Panel A shows the result of a comparative risk analysis conducted on longitudinal data of project status from February 2024-2026. The orange line shows the progression of projects from at-risk status to withdrawal during the first two years following at-risk designation, while the green line shows the same progression but for recovery. Panel B shows the same information except that a group of related projects which recovered from on-hold status were removed because their on-hold designation was not related to an official Verra investigation.*

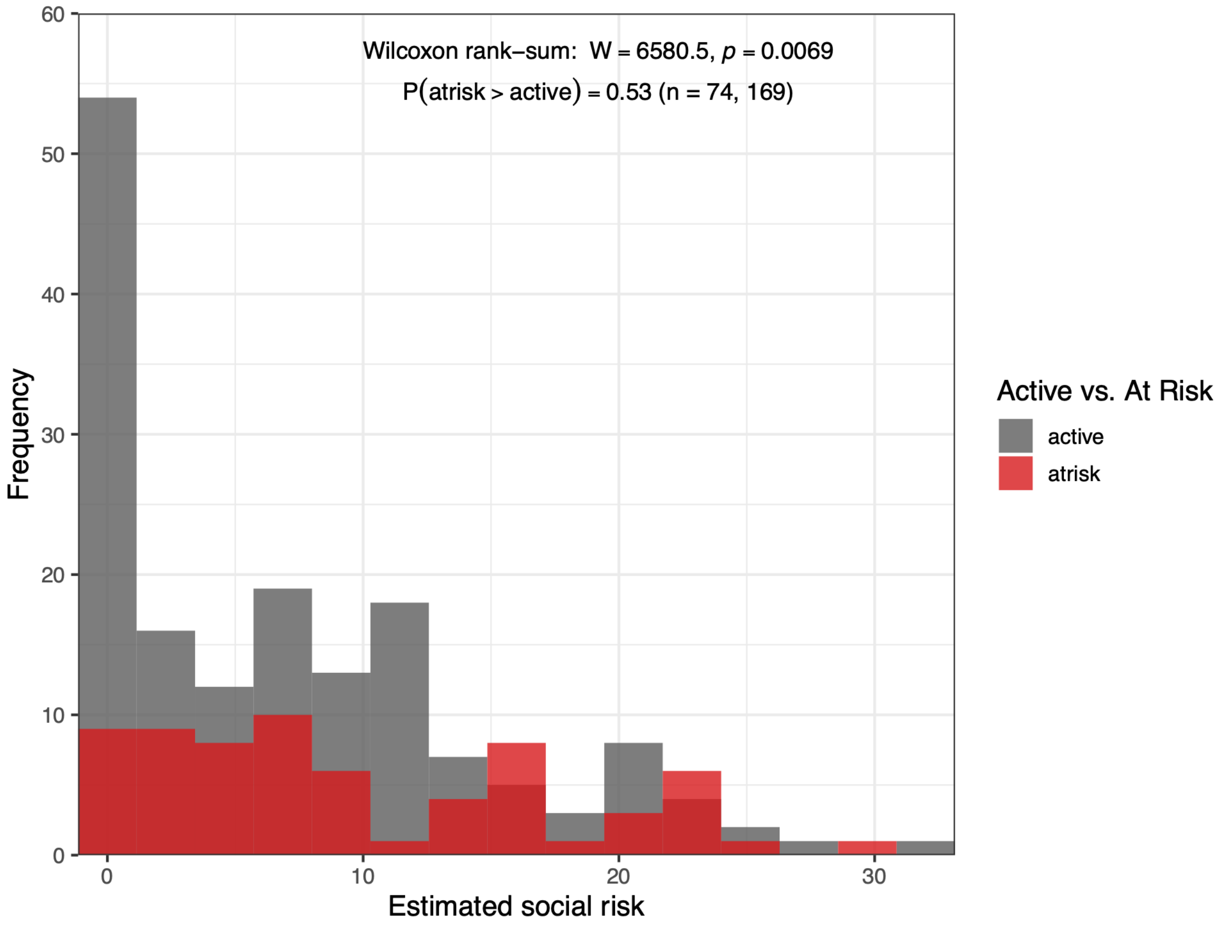


*Figure S4 – Comparison of the distribution of self-assessed social risks between active (gray) and at-risk (red) projects. The results of a Wilcoxon rank-sum test are shown at the top of the panel. Risk is expressed as the percent of issued VCUs that would be lost over a 100-year period.*

**Table S1: At-risk forest carbon offset projects, ranked by estimated annual emissions reductions**

| *Project name* | *Activity type* | *Status* | *Annual emissions reductions ($tCO_2e/yr$)* | *Social risk* | *Natural risk* | *Total risk* |
|---|---|---|---|---|---|---|
| Kariba REDD+ Project | REDD | Withdrawn | 6,550,464 | 12% | 1.25% | 13% |
| Rimba Raya Biodiversity Reserve Project | REDD | Late-to-verify | 3,527,171 | 4% | 3.25% | 12% |
| NIHT Topaiyo REDD+ Project | REDD | Late-to-verify | 2,262,521 | 17% | 2% | 19% |
| Punjab Agroforestry Project II | ARR | On-hold | 1,571,328 | 17% | 2% | 19% |
| Evergreen REDD+ Project | REDD | On-hold | 1,566,547 | 5% | 2% | 12% |
| Makira Forest Protected Area, Madagascar | REDD | Late-to-verify | 1,267,231 | 8% | 5% | 13% |
| Punjab Agroforestry Project I | ARR | On-hold | 1,098,123 | 17% | 2% | 19% |
| CAZ Protected Area Project, Madagascar | REDD | Late-to-verify | 1,011,973 | 4% | 2% | 12% |
| Rio Anapu-Pacaja REDD Project | REDD | On-hold | 938,000 | 10% | 1.75% | 12% |
| Guinea-Bissau REDD Project | REDD | Late-to-verify | 920,436 | 6% | 0% | 12% |
| Liugui Afforestation Project | ARR | Withdrawn | 878,138 | 2% | 0% | 12% |
| Qinghai Afforestation Project | ARR | On-hold | 766,948 | 7% | 1.5% | 12% |
| Qianbei Afforestation Project | ARR | Withdrawn | 708,123 | 2% | 0% | 12% |
| Gansu Dingxi Afforestation Project | ARR | On-hold | 706,810 | 5% | 0.75% | 12% |
| Huadu Afforestation Project | ARR | Withdrawn | 702,405 | 2% | 6% | 12% |

| *Project name* | *Activity type* | *Status* | *Annual emissions reductions (tCO₂e/yr)* | *Social risk* | *Natural risk* | *Total risk* |
|---|---|---|---|---|---|---|
| Chudu Afforestation Project | ARR | Withdrawn | 668,883 | 2% | 0% | 12% |
| Anhuang Afforestation Project | ARR | Withdrawn | 607,596 | 2% | 0% | 12% |
| Haidong Afforestation Project | ARR | On-hold | 598,145 | 7% | 1.5% | 12% |
| RMDLT Portel-Para REDD Project | REDD | On-hold | 553,273 | 10% | 1.75% | 12% |
| Jilin Linjiang Afforestation Project | ARR | On-hold | 546,751 | — | — | — |
| Unitor REDD+ Project | REDD | On-hold | 522,923 | 5% | 4% | 12% |
| Xining City Afforestation Project | ARR | On-hold | 499,982 | 7% | 1.5% | 12% |
| Henan Nanzhao Afforestation Project | ARR | Withdrawn | 493,806 | 0% | 1.25% | 12% |
| Hechu Afforestation Project, Anhui | ARR | Withdrawn | 469,187 | 2% | 0% | 12% |
| Miaoling Afforestation Project | ARR | Withdrawn | 458,449 | 2% | 1.75% | 12% |

*Activity types: ARR = afforestation, reforestation, and restoration; REDD = reduced emissions from deforestation and degradation; IFM = improved forest management. Risk values are self-assessed percentages from each project's non-permanence risk report; social risk is the sum of external and internal risk. "—" indicates a missing value. Status reflects each project's designation in the VCS registry as of February 17, 2026.*

### Table S2: Required social-risk buffer contribution to ensure durability under observed failure rates

| *Model* | *Horizon (years)* | *Required budget (observed rate = 43%)* | *Required budget (observed rate = 68%)* |
|---|---|---|---|
| Weibull | 40 | 42.1% [39.6%, 43.0%]<br>*4.2× current budget, 3.2× total contribution* | 66.5% [62.6%, 67.9%]<br>*6.6× current budget, 5.0× total contribution* |
| Weibull | 100 | 43.0% [43.0%, 43.0%]<br>*4.3× current budget, 3.2× total contribution* | 68.0% [68.0%, 68.0%]<br>*6.7× current budget, 5.1× total contribution* |
| Lognormal | 40 | 38.1% [34.7%, 40.7%]<br>*3.8× current budget, 2.9× total contribution* | 60.3% [54.9%, 64.3%]<br>*6.0× current budget, 4.5× total contribution* |
| Lognormal | 100 | 42.9% [42.4%, 43.0%]<br>*4.2× current budget, 3.2× total contribution* | 67.8% [67.1%, 68.0%]<br>*6.7× current budget, 5.1× total contribution* |
| Gompertz | 40 | 43.0% [42.3%, 43.0%]<br>*4.3× current budget, 3.2× total contribution* | 68.0% [66.9%, 68.0%]<br>*6.7× current budget, 5.1× total contribution* |
| Gompertz | 100 | 43.0% [43.0%, 43.0%]<br>*4.3× current budget, 3.2× total contribution* | 68.0% [68.0%, 68.0%]<br>*6.7× current budget, 5.1× total contribution* |
| Spline | 40 | 34.2% [24.0%, 40.7%]<br>*3.4× current budget, 2.6× total contribution* | 54.1% [38.0%, 64.4%]<br>*5.4× current budget, 4.1× total contribution* |
| Spline | 100 | 42.8% [30.7%, 43.0%]<br>*4.2× current budget, 3.2× total contribution* | 67.7% [48.5%, 68.0%]<br>*6.7× current budget, 5.1× total contribution* |

*Required social-risk budget is calculated as P(at-risk by horizon) × observed at-risk-to-failure rate, for each of our two empirically observed rates: 43% (including the group of related projects placed on hold for minor reporting omissions) and 68% (excluding this group; see Methods and Results, Assessing project failure rates with longitudinal data). Values shown are point estimate [95% CI], with each cell's required budget expressed below as a multiple of the current social-risk budget (10.1% of issued credits: 13.3% average total buffer contribution × 75.6% share allocated to social risk) and of the current average total buffer contribution (13.3%). The 40- and 100-year horizons correspond to VCS's stated monitoring period and reversal risk assessment horizon, respectively. Both horizons lie beyond the maximum follow-up period in our dataset (~25 years); estimates at these horizons require extrapolating fitted hazard models well past the observed data range and should be treated as illustrative rather than precise (see Methods).*
